# Generative AI-based design of hybrid transcriptional activator proteins with new DNA-binding specificity

**DOI:** 10.64898/2025.12.31.697150

**Authors:** Sota L Okuda, Atsushi Minami, Michio Aiko, Kohei Uetsuka, Kazuteru Miyazaki, Kazumasa Ohtake, Daisuke Kiga

**Author notes:** Correspondence to: Daisuke Kiga. These authors contributed equally to this work (Sota L Okuda, Atsushi Minami, Michio Aiko).

## Abstract

Transcriptional control arises from the specific recognition of promoter DNA by transcription factors (TFs), forming the basis of cellular information processing and gene regulation. In synthetic biology, TF-promoter interactions are assembled into genetic circuits to program cellular behaviors. To ensure expected circuit performance, most synthetic genetic circuits rely on combinations of well-characterized and orthogonal transcriptional regulations. This reliance minimizes crosstalk but constrains circuit complexity and information integration. Creating hybrid TFs that combine or interpolate promoter specificities could therefore expand the design space of synthetic regulatory systems. However, it remains unclear whether hybrid functions can be created by mixing amino acid sequences, and how such functional integration could be achieved in a principled manner. Here we show that a variational autoencoder (VAE) trained on LuxR-family DNA-binding domains can generate transcription factors with hybrid and partially novel promoter recognition properties. By sampling intermediate regions of the VAE-learned latent space, we designed hybrid TFs that activate both the *lux* and *las* promoters. High-throughput sort-seq assays together with individual *in vivo* assays revealed that a subset of functional variants exhibited dual-responsive behavior while maintaining sequence-selective DNA recognition. Together, these results provide a data-driven strategy for exploring functional intermediate sequences between closely related proteins.

## Introduction

Transcriptional control, mediated by the precise recognition of promoter DNA by transcription factors (TFs), is fundamental to information processing in natural cells and to the operation of synthetic genetic circuits. In synthetic biology, which aims to engineer biological systems by assembling standardized genetic parts into programmable networks^1^, TF-promoter interactions serve as the fundamental building blocks of synthetic genetic circuits^2,3^. Within such circuits, transcriptional regulation plays a central role, with TFs acting as molecular switches that convert environmental or intracellular cues into gene expression outputs. To simplify design and minimize crosstalk, well-characterized and orthogonal genetic parts have been predominantly used for constructing synthetic genetic circuits^4–10^. While this strategy supports reliable circuit behavior, it also constrains circuit complexity and information density. To construct more sophisticated biological computation, increasingly complex information-processing architectures have been developed across molecular and cellular scales.

Recently, neural network-like pattern recognition and decision-making have been demonstrated using multi-input and multi-output genetic circuits *in vitro*^11–13^ and in living cells^14–21^. Yet most of these systems still depend on assembling multiple components each of which has its major target, rather than using regulatory elements that combine multiple input/output specificities within a single element. Therefore, enabling compact genetic circuits with multi-input and multi-output transcriptional logic requires transcription factors and promoters that interact across multiple regulatory axes. One promising strategy to achieve this is the design of dual-active hybrid TFs that combine recognition features of multiple parental regulators (**Fig. 1a**).

**Fig. 1.**
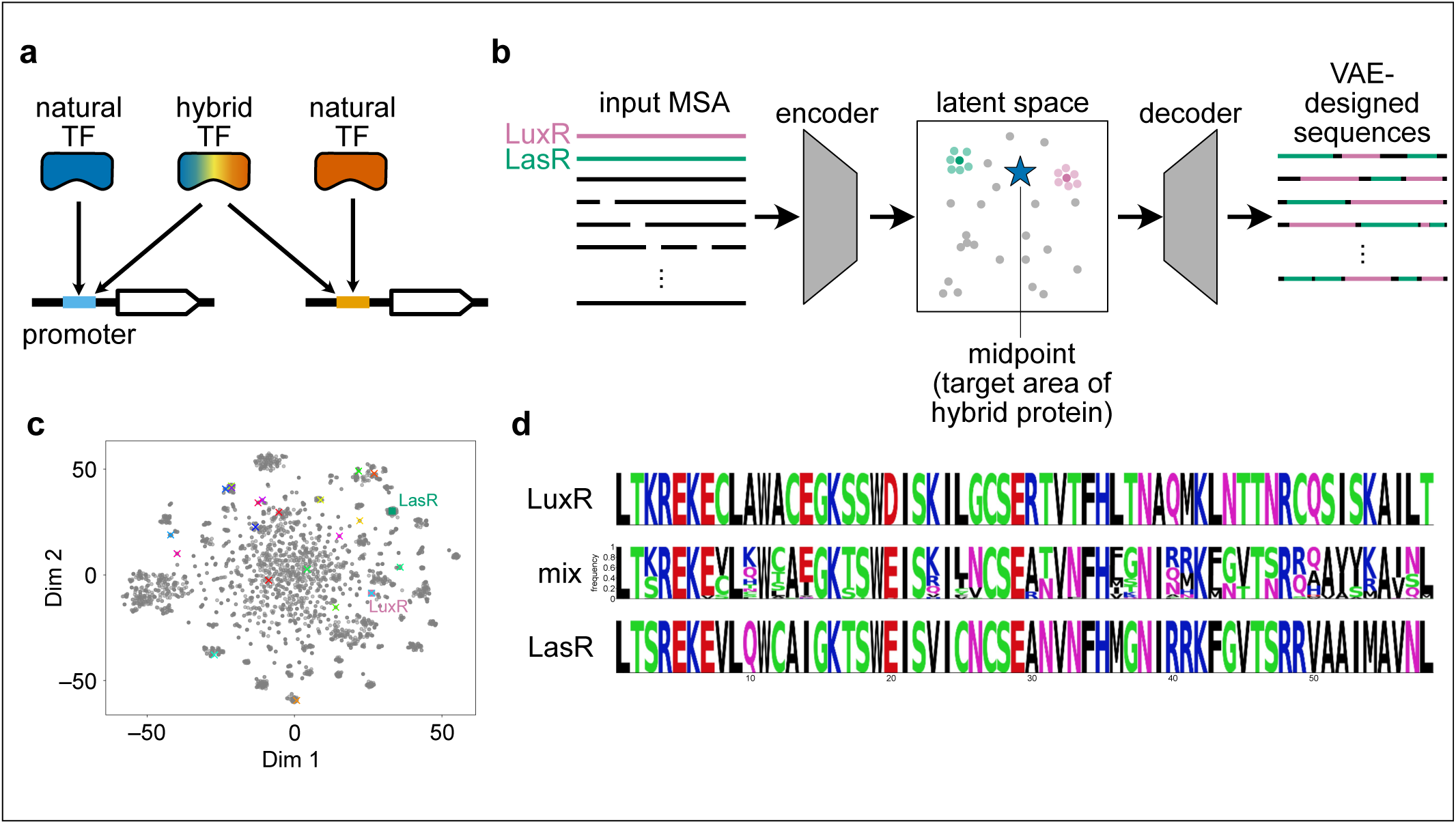
Schematic of the VAE-based sequence generation strategy for hybrid TFs. **a,** Design and dual-sequence recognition of the hybrid transcription factor. Parental natural transcription factors (natural TFs: blue and orange) specifically bind to their respective target DNA sequences. This study aims to design the hybrid TF (colored in rainbow), formed by mixing of natural TFs, that exhibits dual recognition capability, enabling it to bind to both sequence and activate downstream gene expression for both targets. **b,** Schematic overview of the MSA-VAE architecture used for hybrid sequence design. Aligned LuxR-family DNA-binding domains (DBDs) are encoded into a low-dimensional latent representation and decoded back into sequence space. **c,** t-SNE projection of representative LuxR-family transcription factors. The positions of LuxR and LasR are indicated by a pink square and a green square, respectively. Additional representative family members are plotted using X-shaped markers. Natural sequences are shown as individual points. **d,** Sequence sampling from the LuxR-LasR interpolation region. Hybrid sequences were generated from a hyperspherical region centered at the midpoint between the LuxR and LasR latent vectors (0.5-0.5 interpolation), decoded, and filtered for redundancy using CD-HIT.

Given that protein function is ultimately dictated by its amino acid sequence, including DNA recognition by TFs, could hybrid sequences give rise to hybrid functions? Several approaches have been explored to address this question, including recombination-based methods such as domain swapping^22^ and DNA shuffling^23^. Ancestral sequence reconstruction (ASR)^24^ offers a conceptually distinct route, inferring intermediate sequences by tracing evolutionary trajectories within a protein family. These methods have enabled the creation of chimeric proteins or functional intermediates between homologous proteins. However, these approaches do not account for the residue-residue interactions and sequence context that collectively determine folding, stability, and DNA-recognition specificity in TFs.

Recent advances in deep generative modeling offer a fundamentally different way to mix sequence information. Variational autoencoders (VAEs) are well known from image generation for their ability to interpolate smoothly between examples, such as producing intermediate handwritten digits by sampling along continuous trajectories in latent space^25^. VAEs have recently been applied to protein sequence design, where VAEs trained on natural sequence families can learn evolutionary constraints and generate functional variants of enzymes, signal peptides, and nanobodies without requiring structural information^26–33^. Notably, however, these applications have focused on producing alternative variants within a single functional class; whether latent-space interpolation can generate meaningful “intermediate” sequences that combine functional features of two homologous transcription factors remains to be elucidated.

In this study, we employed a VAE to generate hybrid transcription factor sequences derived from the parental TFs; LuxR and LasR. By constructing a diverse library of VAE-generated proteins and assessing their transcriptional activation abilities, we identified several proteins exhibiting dual promoter-recognition capabilities.

Furthermore, by combining a randomized promoter library with deep sequencing (sort-seq), we systematically profiled the promoter recognition specificities of each variant. Together with structural analyses based on AlphaFold3 predictions and molecular dynamics simulations, we reveal the molecular basis underlying the DNA-binding properties of these hybrid proteins. Our results provide valuable insights into the rational design of proteins with hybrid functions.

## Results

### Design of hybrid transcription factors using a variational autoencoder (VAE)

We first sought to design hybrid transcription factors by applying a variational autoencoder (VAE) to the DNA-binding domains (DBDs) of LuxR-family regulators. LuxR-type transcription factors consist of an N-terminal ligand-binding receiver domain and a C-terminal helix-turn-helix DBD that mediates promoter recognition^34^. Since the DBD alone is sufficient to bind target DNA and can activate transcription independently of ligand binding^35,36^, we focused our design efforts exclusively on this domain. This allowed us to directly probe how sequence-level changes within the DBD alter promoter specificity.

To generate hybrid sequences, we constructed a curated multiple sequence alignment (MSA) of LuxR-family DBDs^37^ and trained an MSA-VAE on this dataset^27,29,33^. The VAE comprises an encoder that compresses each aligned sequence into a low-dimensional latent representation and a decoder that reconstructs sequences from these representations. During training, the model simultaneously optimizes a reconstruction objective and a regularization term that encourages the latent variables to follow a multivariate Gaussian distribution, resulting in a structured latent space that captures biophysical constraints in the input sequences (**Fig. 1b**). When encoded by the trained model, the parental LuxR and LasR sequences occupied distinct regions of the latent space (**Fig. 1c**). Other representative members of the LuxR family, including RhlR and CinR, were also mapped to distinct positions, with clusters of related natural sequences forming around each representative point (**Fig. 1c**). This segmentation indicates that the diversity present in the training dataset is faithfully preserved in the latent space and that the VAE captures family-level sequence relationships rather than collapsing them into a single region. The clustering among these subfamilies also enabled the definition of an interpolation point between the LuxR and LasR latent vectors, providing a principled way to explore sequence variants that combine features of both regulators.

Using this latent-space representation, we sampled sequences from a hyperspherical region centered at the midpoint between the LuxR and LasR latent vectors (0.5-0.5 interpolation). This strategy enabled targeted sampling of hybrid sequences, in contrast to prior VAE studies that sampled amino acid sequences closely related or neighboring locations in the latent space for collecting similar variants^27,29,32,33^. Sampling across the interpolation region produced a diverse set of candidate sequences that contain novel combinations of parental residues while retaining global amino-acid usage patterns of the LuxR family. From each of the top-performing VAE models, 20,000 sequences were decoded from the local interpolation region. After removing redundant and gap-containing sequences using CD-HIT, we obtained a pool of unique candidate hybrids (**Fig.1d**).

### Functional characterization of VAE-designed TFs toward lux and las promoters

To determine whether the VAE-designed hybrid sequences exhibit functional activity *in vivo*, we selected 9 candidates at random from the generated sequences and evaluated their ability to activate transcription from the native *lux* and *las* promoters in *Escherichia coli*. For each designed variant, we constructed two coding sequences that differed only in their immediate N-terminal context: one beginning with Met (“M+CDS”), and the other beginning Met followed by Ala (“MA+CDS”) to account for potential N-terminal context effects^38,39^, resulting in a total of 18 constructs. The mixed oligonucleotide pool was cloned and expressed in *E. coli*, and its ability to activate transcription from the native *lux* and *las* promoters was quantified using GFP reporter assays (**Fig. 2a**). Some of individual fluorescent colonies were isolated, and the corresponding coding sequences were identified by sequencing prior to quantitative analysis. Variants that were not recovered for quantitative analysis likely reflect a combination of factors, including severe growth defects, unstable or insufficient expression, lack of detectable promoter activation, or sampling bias inherent to colony-based isolation. The impact of the N-terminal Ala insertion was heterogeneous across variants, with no consistent trend observed between the two forms.

**Fig. 2.**
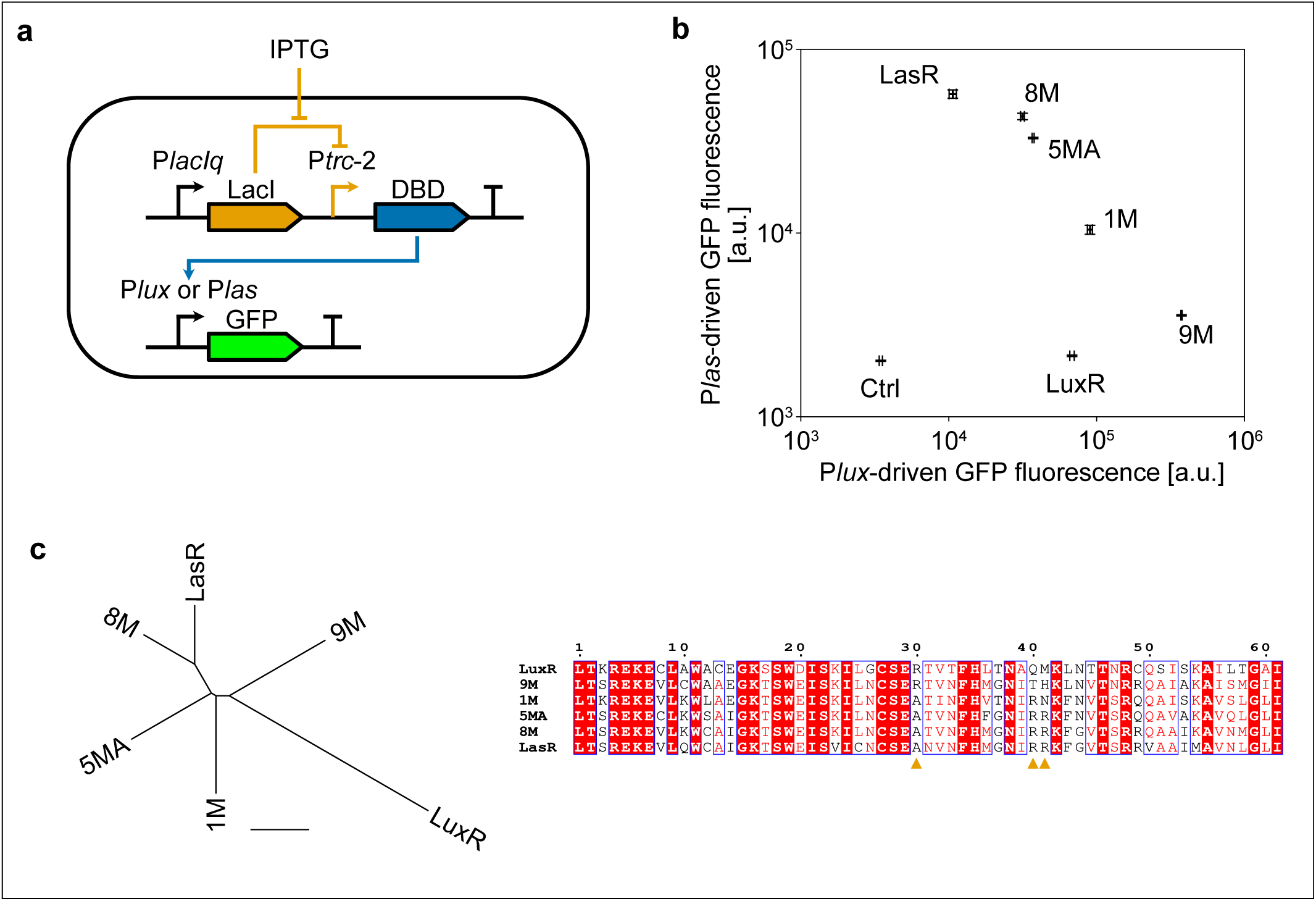
Functional validation of VAE-generated hybrid TFs. **a,** Schematic overview of the GFP reporter assay used to quantify transcriptional activation by VAE-designed transcription factors. Each construct was expressed in *E. coli*, and promoter activity was measured from GFP driven by the native *lux* or *las* promoter. **b,** Transcriptional activation activities of VAE-designed variants and wild-type LuxR and LasR toward the *lux* and *las* promoters. Bars represent the geometric mean of GFP fluorescence measured by a flow-cytometry. A strain harboring pET16b (an empty vector) is shown as a control (Ctrl). Data are shown as the mean of three biological replicates, and error bars indicate the standard error of the mean. **c,** (left) Unrooted phylogenetic tree of the parental LuxR-family proteins and the four dual-responsive variants (8M, 5MA, 1M, and 9M). The tree was inferred from the LuxR-family DBD alignment and visualized without an assigned root. (right) Multiple sequence alignment of the variants shown in the phylogenetic tree together with the wild-type LuxR and LasR DBDs. Positions corresponding to residues 30, 40, and 41 discussed in the main text are highlighted with orange triangles.

The quantitative assay using a flow-cytometry exhibited a wide range of transcriptional activities for each TF across the two promoters (**Fig. 2b**). Under identical assay conditions, wild-type LuxR showed robust activation of the *lux* promoter but negligible activity at the *las* promoter, whereas wild-type LasR activated the *las* promoter strongly and displayed weak but detectable activity toward the *lux* promoter. Notably, a subset of constructs activated both promoters to varying degrees, revealing balanced dual-responsive behavior that is not observed for the parental LuxR or LasR proteins. Among the variants that activated both promoters, four constructs (8M, 5MA, 1M, and 9M) displayed distinct activity profiles. Variants 8M and 5MA showed higher activation at the *las* promoter than at the *lux* promoter, whereas 9M exhibited the opposite trend and 1M showed an intermediate balance between the two parental proteins (**Fig. 2b**). Consistent with these functional differences, an unrooted phylogenetic analysis placed 8M and 5MA closer to LasR, while 9M was positioned nearer to LuxR, with 1M occupying an intermediate branch (**Fig. 2c**). Sequence alignment of these variants further revealed residue-level features that parallel their activity patterns: 9M uniquely retained Arg30 conserved in LuxR subfamily, whereas the other three variants carried the Ala at this position conserved in LasR subfamily; and both 8M and 5MA preserved LasR-associated Arg40 and Arg41 (**Supplementary Fig. S1**). Together, these observations indicate that promoter preference in the VAE-designed hybrids reflects combinations of parental sequence features at key DNA-contacting positions.

### Promoter recognition profiles of VAE-designed proteins revealed by sort-seq

Building on the initial characterization of clones from 18 variants pool, we next assessed promoter recognition across a larger set of VAE-designed sequences using a massively parallel reporter assay (MPRA)^40,41^. Sixty additional VAE-designed DBD sequences were randomly selected from the VAE-generated candidate sequences, and two versions of each were created: one with M+CDS and the other with MA+CDS, yielding a total of 120 unique variants. These sequences were synthesized as an oligonucleotide pool and collectively cloned into the expression vector to construct a pooled plasmid library, which was then transformed into *E. coli* harboring a GFP reporter plasmid under the control of either *lux* or *las* promoter. Fluorescence from each cell was measured by a fluorescence-activated cell sorter (FACS), and cells were sorted into predefined fluorescence bins for each promoter. Plasmid DNAs from each bin were extracted and subjected to deep-sequencing (sort-seq)^42,43^, with activity of each variant quantified by counting exact matches to the designed sequences across bins, normalizing these counts by the fraction of cells in each bin of histogram from FACS analysis, and calculating a geometric mean activity score for each promoter (**Fig. 3a; Supplementary Fig. S2**).

**Fig. 3.**
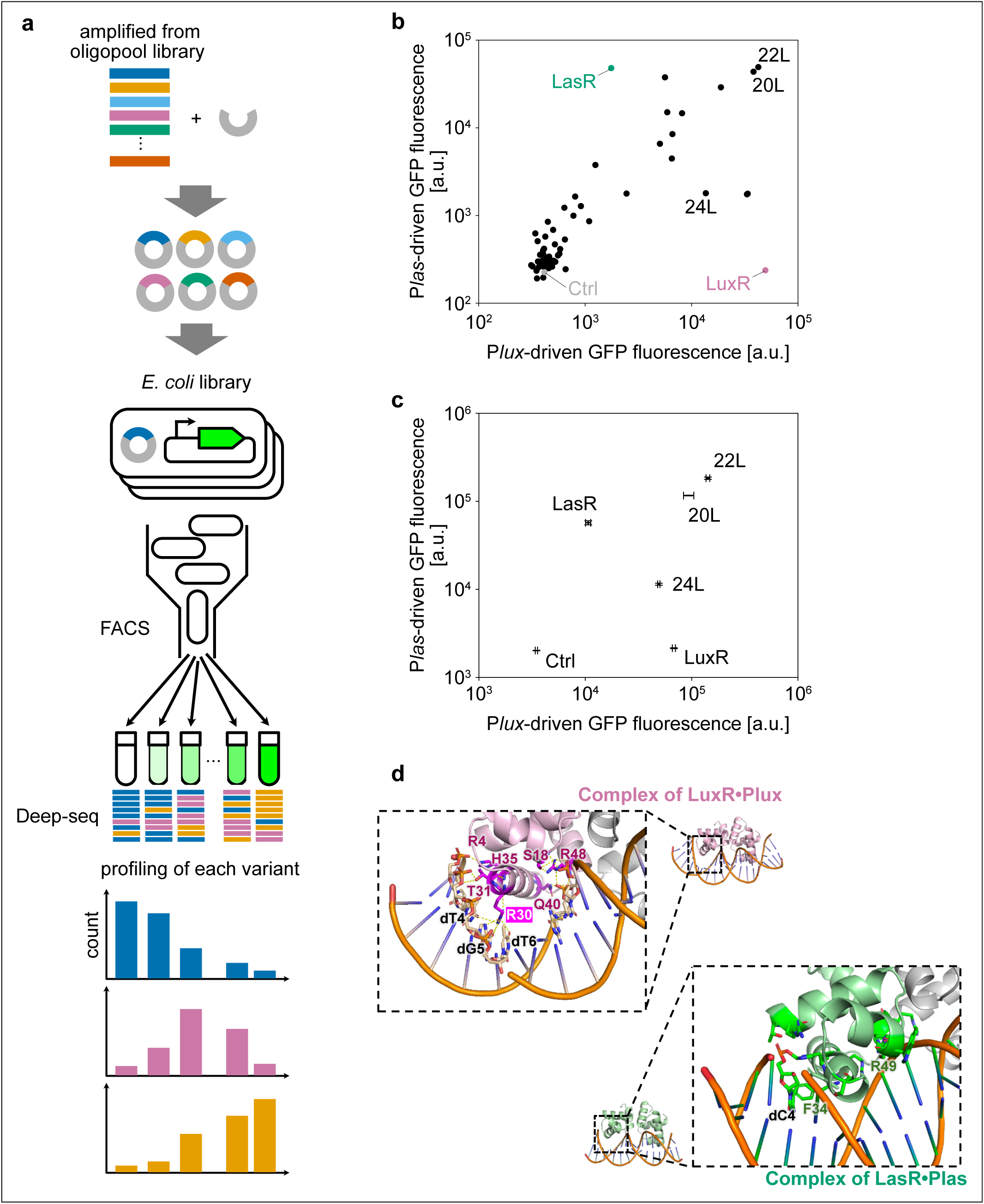
Massively parallel reporter assays and structural features of VAE-designed TFs. **a,** Schematic overview of the sort-seq workflow. A total of 120 VAE-designed variants were constructed as a pooled plasmid library, transformed into *E. coli*, and sorted into fluorescence bins based on reporter output from the *lux* or *las* promoter. Plasmid DNA recovered from each bin was deep-sequenced, and activity scores were calculated for each variant from exact sequence counts across bins. **b,** Promoter-activation profiles of the 120 VAE-designed variants obtained from one of two biological replicate sort-seq experiments. The result of the replicate is shown in **Supplementary Fig. S4**. Each point represents the geometric mean activity score of a variant toward the *lux* or *las* promoter. Variants showing measurable activation include lux-biased, las-biased, and dual-responsive sequences. **c,** Validation of pooled-screen results by individual clone assays. Variants 20, 22, and 24 (20L, 22L, and 24L) were cloned individually and assessed using the GFP reporter assay. The individually measured activities recapitulate the promoter-recognition patterns inferred from the pooled deep-sequencing data. **d,** Representative structural models of parental transcription factors bound to their cognate promoters. AlphaFold-based models of LuxR bound to the *lux* promoter (left) and LasR bound to the *las* promoter (right) after molecular dynamics refinement. Proteins are shown in cartoon representation with selected DNA-contacting residues highlighted as sticks, and DNA is shown as a double helix.

Sixteen variants of the 120 VAE-designed proteins in the sort-seq exhibited measurable activation toward either promoter. Notably, many of these active variants showed detectable responses at both promoters, indicating that dual-responsiveness was a frequent outcome within the functional subset (**Fig. 3b**). These variants displayed a broad dynamic range of activities, spanning strongly *lux*-biased, strongly *las*-biased, and dual-responsive profiles. Despite this diversity, dual-responsive variants were more prevalent than strictly promoter-specific ones. Overall, the population-level trends captured by the sort-seq reflected the behaviors identified in the single-construct assays, supporting the robustness of the functional patterns observed.

To validate the activities inferred from the sort-seq, we individually constructed plasmids encoding selected variants (No. 20, 22, and 24, designated as 20L, 22L, and 24L, respectively) and measured their promoter activation using the GFP reporter assay described above (**Fig. 2a**). The individually tested clones reproduced the activity patterns observed in the pooled screen, confirming the dual-responsive behaviors inferred from the sort-seq data (**Fig. 3c**). These results demonstrate that the sort-seq reliably captures the promoter-recognition properties of VAE-designed transcription factors.

### Structural insights into promoter recognition by the transcription factors

To understand how LuxR-family transcription factors recognize their cognate promoters and to identify residues potentially responsible for the altered specificity observed in the VAE-designed variants, we modeled the complexes between LuxR or LasR and the *lux* or *las* promoter sequences using AlphaFold3^44^. Four protein-DNA complexes (LuxR•P*lux*, LuxR•P*las*, LasR•P*lux*, and LasR•P*las*) were initially predicted, and the resulting structures were subjected to molecular dynamics simulations to examine their stability and residue-level interactions.

Although global structural metrics such as RMSD and overall binding energetics did not fully distinguish the four complexes (**Supplementary Fig. S3a**), the simulations revealed reproducible patterns in specific residues that contribute to nucleotide recognition in each protein (**Fig. 3d**). In LuxR, Arg30, described above, formed the only consistently stable hydrogen bonds with specific bases (dT4, dG5, and dT6) within the major groove, identifying it as a primary determinant of LuxR-specific base recognition.

This position is Ala30 in LasR, explaining the weaker and less specific base contacts predicted for LasR at the same sites. In contrast, LuxR Thr31 engaged the DNA phosphate backbone, suggesting a role in stabilizing DNA positioning rather than dictating base identity; LasR carries an Asn31 at this site, which may alter backbone interactions without imposing strong base preferences. A conserved Phe34 in both LuxR and LasR was positioned for potential T-shaped π-π stacking with DNA bases (**Supplementary Fig. S3b**), consistent with analogous interactions reported in another transcription factor MarA^45^. LasR also contained two positively charged residues, Arg40 and Arg49, located near the major groove; although direct hydrogen bonds were not consistently observed in MD trajectories, both residues contributed strongly to total DNA-binding energy, whereas the corresponding LuxR residues (Gln40 and Cys49) provided less energetic stabilization. These differences highlight distinct recognition strategies employed by the two parental regulators.

Notably, the VAE-designed variants 20L and 22L combined features of both parental modes of DNA engagement (**Supplementary Fig. S3c**). At the Arg/Ala30 position, both variants carried the LasR-type Ala30, consistent with reduced LuxR-specific base contacts. However, they also preserved the LuxR-type Thr31, potentially maintaining similar DNA-backbone interactions. The LasR-associated Arg40 was retained in both variants, while position 49 was occupied by Gln, intermediate in polarity between the LuxR and LasR residues (Cys and Arg, respectively), suggesting partial recapitulation of LasR-like electrostatic stabilization. Together with the conservation of Phe34 and the universally retained Arg at position 48, these patterns suggest that 20L and 22L adopt hybrid combinations of the parental DNA-contacting residues, providing a structural basis for their mixed and broadened promoter-recognition profiles.

### Promoter recognition profiles of VAE-designed proteins revealed by deep sequencing

Since the VAE-designed proteins exhibited broadened activities at the native *lux* and *las* promoters, it was essential to determine whether their apparent dual responsiveness reflected genuine sequence preferences or merely nonspecific DNA binding. To address this, we generated randomized promoter libraries by mutating nucleotide positions presumed to influence recognition by LuxR-family regulators and then assessed promoter-recognition specificity for wild-type LuxR, wild-type LasR, and VAE-designed selected variants (20L and 22L) using an MPRA. The randomized promoter libraries were synthesized from random oligonucleotides and cloned upstream of a GFP reporter to create pooled plasmid libraries. Either wild-type LuxR, wild-type LasR, 20L, or 22L was expressed in *E. coli* containing the promoter library, and fluorescence output from individual cells was quantified by FACS (**Fig. 4a**). Cells were sorted into multiple fluorescence bins for each transcription factor, and promoter sequences from each bin were recovered and deep-sequenced (sort-seq). For each promoter variant, we calculated an enrichment-based activity score from its distribution across fluorescence bins.

**Fig. 4.**
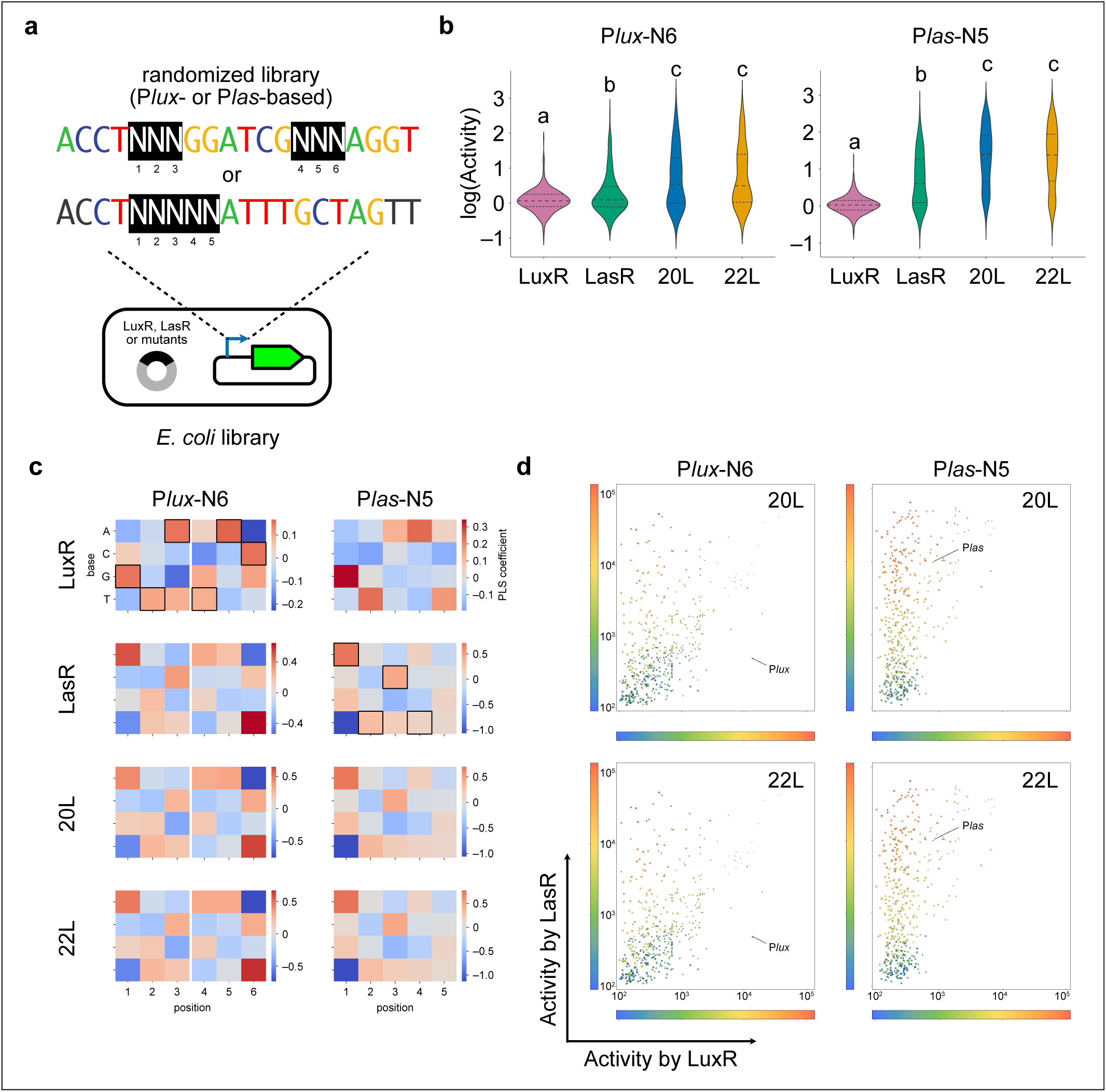
Specificity of VAE-designed transcription factors toward randomized promoter libraries. **a,** Schematic of the promoter library construction used for sort-seq. Randomized promoter libraries were generated by mutating selected nucleotide positions within the *lux* and *las* promoter regions and cloned upstream of a GFP reporter. Cells expressing wild-type LuxR, wild-type LasR, or VAE-designed variants (20L or 22L) were sorted into multiple fluorescence bins by FACS, and promoter sequences from each bin were recovered and analyzed by deep sequencing to calculate enrichment-based activity scores. **b,** Violin plots showing the distribution of activity scores across all randomized promoter variants for each transcription factor. Each violin illustrates the full activity distribution for wild-type LuxR, wild-type LasR, 20L, or 22L. Horizontal bars indicate group means. Pairwise differences in mean activity among the four DBDs were assessed by one-way ANOVA followed by post hoc multiple-comparison testing. **c,** Nucleotide-enrichment profiles derived from the randomized promoter libraries. Position-wise nucleotide preferences are shown for wild-type LuxR, wild-type LasR, 20L and 22L. The nucleotide identities of the native *lux* and *las* promoters are indicated by black outlines at positions previously reported to be conserved based on sequence logo analyses^46,47^, shown in the corresponding LuxR and LasR heatmaps, respectively. **d,** Four-dimensional activity landscape of promoter responses to wild-type and VAE-designed transcription factors. Promoter activities for wild-type LasR and LuxR are plotted on the horizontal and vertical axes, respectively. Activities for 20L or 22L are encoded by color, and negative-control fluorescence is represented by point size.

Analysis of the randomized promoter libraries revealed that the VAE-designed variants 20L and 22L did not simply exhibit indiscriminate DNA binding. Although the VAE-designed variants showed higher mean activities across the promoter library, their activity distributions exhibited greater variance than those of the wild-type proteins. This broader spread reflects an expanded dynamic range of promoter activation rather than loss of sequence-selective recognition (**Fig. 4b**). To further examine how their sequence preferences were altered, we quantified nucleotide-level enrichment across randomized positions for each transcription factor (**Fig. 4c**). Both 20L and 22L displayed preference profiles broadly similar to those of LasR, especially in the *las* promoter library. These trends align with structural modeling, which suggested that LasR makes broader and less restrictive base contacts than LuxR, providing a plausible route by which LasR-like variants could acquire activity at the *lux* promoter (**Fig. 3c, d**). In contrast, analysis of the *lux* promoter library revealed that LuxR-like nucleotide preferences were partially retained. For example, at position 2, LasR favors G over T, whereas LuxR most strongly prefers T; notably, 20L and 22L similarly exhibited a T > G preference at this position. At position 1, both 20L and 22L retained a primary preference for A, similar to LasR, while additionally exhibiting increased tolerance for G, which is most preferred by LuxR. The nucleotide preference patterns observed for wild-type LuxR and LasR were consistent with those reported in previous studies, supporting the validity of the assay^46,47^. Notably, furthermore, the VAE variants preferentially activated a subset of sequences not strongly activated by both wild-type TF, as revealed by mapping promoter activities into a four-dimensional activity landscape where LuxR and LasR activities are plotted on two axes and the 20L or 22L activity is represented by point coloration (**Fig. 4d**). This pattern indicates that the VAE-designed factors do not simply interpolate between LuxR and LasR specificity but instead acquire hybrid or partially novel recognition profiles.

## Discussion

Our study demonstrates that a variational autoencoder can effectively design hybrid transcription factors that bridge functional features of distinct LuxR-family regulators. By training an MSA-VAE on a curated alignment of LuxR-family DNA-binding domains and sampling sequences from the interpolated region between LuxR and LasR in latent space, we generated a series of hybrid DBDs that exhibited functional promoter activation *in vivo*. Notably, several of these VAE-designed proteins showed dual-responsive activity toward both the *lux* and *las* promoters, which is not naturally observed in either parental protein. This finding provides direct experimental support for the idea that intermediate sequences between homologous transcription factors can manifest intermediate or even expanded promoter-recognition capacities.

Conventional strategies for combining functionalities from homologous transcription factors such as domain swapping or DNA shuffling are inherently limited in how finely they can mix parental features. Because these approaches recombine blocks of residues rather than individual amino acids, they often disrupt local residue-residue and residue-nucleotide interactions that are essential for DNA recognition. Ancestral sequence reconstruction (ASR) offers an alternative, evolutionarily grounded route toward “intermediate” sequences, but in our experiments the ASR-derived candidates showed LuxR-like behavior: one reconstructed sequence expressed poorly, and the remaining two activated only the *lux* promoter (**Supplementary Fig. S5**). This limited divergence may reflect the intrinsic constraints of ASR, which infers residues independently at each position and therefore cannot capture co-variation of residues or inter-residue interactions that support stable DNA-binding interfaces. Moreover, ASR reconstructs the most probable ancestral states given the multiple sequence alignment; consequently, biases in the underlying sequence dataset may be propagated into the reconstructed sequences.

Beyond classical approaches, emerging protein design technologies also offer alternative routes for engineering of transcription factors. For example, large protein language models, trained on millions of diverse sequences, generate latent spaces that capture broad biochemical trends^48,49^. However, it may not resolve local functional distinctions among closely related regulators such as LuxR and LasR. Interpolation in such global resolution is therefore unlikely to yield meaningful mixtures of parental recognition features without additional constraints or task-specific fine-tuning. Additionally, *de novo* DNA-binding domain design^50^ enables the creation of entirely new HTH scaffolds with programmable nucleotide preferences. As *de novo* frameworks continue to improve, the design of hybrid or multi-specific transcription factors may become achievable, providing an exciting future direction for synthetic genetic circuit engineering.

Structural modeling and MD simulations provided a mechanistic basis for the promoter-recognition behaviors observed in the VAE-designed transcription factors. LuxR and LasR differ at several DNA-contacting residues that shape their respective base-recognition modes. LuxR forms more restrictive base-specific interactions, and LasR rather relies on a broader and more permissive contact network (**Fig. 3d**). These structural trends align with the sort-seq promoter library data: the VAE-designed variants (20L and 22L) exhibited LasR-like nucleotide preferences while also activating a subset of promoter sequences not efficiently recognized by either wild-type factor. Visualization of promoter activities in a multidimensional activity landscape further revealed that these engineered proteins occupy promoter-recognition regimes intermediate to, yet distinct from, both parents. Together, the structural and sort-seq results suggest that the VAE-generated hybrids achieve dual responsiveness not through indiscriminate DNA binding, but by combining permissive LasR-like contacts with specific contributions from individual residues, enabling access to a partially novel specificity space.

While this study demonstrates that a VAE trained on natural LuxR-family sequences can generate transcription factors with hybrid or partially novel promoter specificities, several limitations remain. Our experimental validation focused exclusively on the LuxR-LasR axis, and the generality of this framework across other transcription factor families remains to be tested. In addition, VAE-based design relies on the evolutionary diversity present in the underlying sequence alignment; families with sparse sequence representation may prove more challenging. Despite these constraints, extending this framework to larger families, integrating richer functional readouts, and iteratively updating the model with experimental data will further enhance its capacity to design proteins such as transcriptional regulators for increasingly complex synthetic genetic circuits.

## Methods

### Creation of a dataset of LuxR-family proteins

LuxR-family protein sequences were collected based on the LuxI/R dataset reported by Miranda *et al*^37^. The representative LuxI/R query sequences used in that study, together with the LuxR and LasR sequences (NCBI RefSeq accession ID: WP_213090604.1 and WP_003082999.1, respectively), were added to the dataset and aligned using MAFFT^51^. Because all sequences in the dataset are arranged as LuxI (N-terminal) followed by LuxR (C-terminal), the boundary between the LuxI and LuxR regions was manually determined by referencing conserved N-terminal residues among the representative receiver proteins. The LuxI region was then removed to retain only the LuxR-family portion. Sequences that became excessively short after gap removal (shorter than 200 aa), contained large internal gaps, or were duplicates were iteratively filtered out using CD-HIT^52^. MAFFT alignment and filtering were repeated for several rounds to obtain a consistent LuxR-family multiple sequence alignment.

The DNA-binding domain (DBD) dataset was derived from the LuxR-family alignment using experimentally validated N-terminal deletion boundaries of the LuxR and LasR variants used in our laboratory. These representative sequences were added to the dataset, realigned with MAFFT, and the conserved leucine residue marking the N-terminal DBD boundary was identified. In this study, this conserved leucine was defined as residue 1 for all subsequent numbering. Regions upstream of the aligned leucine position were removed, and C-terminal trimming was performed to match the boundaries used by Miranda *et al*. Sequences shorter than 56 aa after trimming or containing extensive gaps were removed, and identical sequences were clustered using CD-HIT. MAFFT alignment and gap-based filtering were repeated twice to increase alignment consistency and information density. The resulting dataset contained only the DBD regions of LuxR-family proteins.

### Model building and validation using VAE

We implemented a multiple-sequence-alignment variational autoencoder (MSA-VAE) in PyTorch following architectures described by previous studies^29,33^. The model was trained to maximize the evidence lower bound (ELBO), consisting of a binary cross-entropy (BCE) reconstruction term and a Kullback-Leibler divergence (KLD) regularization term:

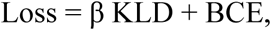

where β is a positive weighting coefficient. The KLD term encourages the encoder posterior *q*(*z*∣*x*) to remain close to a standard normal prior, thereby promoting a smooth and generalizable latent space. BCE was computed per aligned position using one-hot encoded amino-acid categories^25^.

To identify suitable model configurations, we performed a two-stage grid search over (i) the dimension of the latent space, (ii) the size of hidden layers, (iii) the β coefficient for the KLD term, and (iv) random seeds (For detail, see **Supplementary Methods**). Models were trained with a batch size of 32, a learning rate of 1×10^−3^, and the Adam optimizer. Early stopping was applied when the validation ELBO failed to improve for 5 consecutive epochs.

Model quality was assessed using the pairwise amino-acid frequency correlation between generated sequences and the natural sequences in the training dataset, as previously described^29^. Higher correlation values indicate that the model captures evolutionary constraints, including covariation patterns. Unlike previous studies that sample uniformly across the latent space, we generated sequences from interpolation regions between two latent vectors corresponding to selected parental sequences^33,37^. Because models with similar hyperparameters can nonetheless yield distinct latent-space geometries, we selected the top 10 models ranked by the evaluation metric and used all of them for sequence generation (**Supplementary Table S1**). For each model, the LuxR and LasR sequences were encoded to obtain their latent means *z*_LuxR_ and *z*_LasR_. A latent interpolation vector was then defined as:

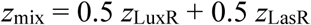

Sequences were sampled from a hyperspherical region centered at *z*_mix_ with a radius *R* was defined as:

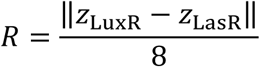

Sampling was performed uniformly within this region by drawing random directions and radii, followed by decoding the resulting latent points into amino acid sequences using the VAE decoder. From this distribution, 20,000 sequences were generated per model. After removing redundant sequences using CD-HIT^52^, nine sequences were randomly selected and used for experimental characterization.

### Structural prediction and MD simulations

The structural models of the parental transcription factors (LuxR and LasR) in complex with *lux* or *las* promoter sequences were predicted using AlphaFold3^44^. Since LuxR-family regulators function as homodimers, all predictions were performed in the dimeric state with the corresponding DNA duplex included as input.

MD simulations were performed using GROMACS 2023.2 with the AMBER99SB-ILDN force field^53,54^ as described previously^55^. Each complex was solvated using the SPC216 water model in a cubic box and neutralized with 150 mM NaCl. Electrostatic and van der Waals interactions were truncated at 1.0 nm, and long-range electrostatics were calculated with the particle mesh Ewald method^56^. Energy minimization was performed using the steepest descent algorithm, and the system was then equilibrated at 300 K and 1 bar for 100 ps each in two steps: first under constant volume (NVT), then under constant pressure (NPT). Temperature and pressure were controlled using the V-rescale thermostat and Parrinello-Rahman barostat, respectively^57^. The LINCS algorithm was used to constrain bonds involving hydrogen atoms^58^. The integration time step was set as 2 fs. Production runs of 100 ns were conducted under the NPT ensemble without restraints, with three independent replicates for each complex. The RMSD and hydrogen bonds profiles were generated by GROMACS modules. Images were generated by PyMOL (The PyMOL Molecular Graphics System, Version 3.1.0 Schrödinger, LLC).

### Construction of plasmids

Plasmids and primers used in this study were listed in **Supplementary Table S2 and S3**, respectively. An expression vector for LuxR DBD was constructed from pPtet LuxR (Ptet-LuxR on pSB3K3)^59^. First, the P*tet*-*luxR* region fragment was excised using EcoRI and PstI and subcloned into the corresponding restriction sites of pSB6A1. Next, inverse PCR was performed to delete the N-terminal region of LuxR (amino acid residues 2-182), followed by self-ligation of the PCR product. An expression vector for the LasR DBD was constructed using oligonucleotides synthesized as an oligo pool (Twist Bioscience, United States). Briefly, double-stranded DNA was generated by primer extension with a complementary oligonucleotide primer, purified, and digested with NheI and HindIII. The resulting fragment was cloned into the corresponding restriction sites of pTrc99A-DsRedExpress2 (for detailed procedures, see **Supplementary Methods**).

For VAE-designed proteins, coding sequences were selected at random from the model-generated sequence pool, codon usage of each sequence is optimized, and a ribosome-binding site was individually designed using the RBS Calculator (v2.1)^60^ to achieve comparable predicted translation initiation rates across variants. Resultant sequences were synthesized as an oligonucleotide library (Twist Bioscience, United States). Double-stranded DNA was generated by primer extension with a complementary oligonucleotide primer, purified, and inserted between the NheI and HindIII sites of pTrc99A-DsRedExpress2, as described above.

Promoter-reporter plasmids were constructed as follows. The gene fragment containing the *las* promoter followed by *gfp* (iGEM Parts Registry: BBa_K649001) was digested with XbaI and SpeI and cloned into the corresponding restriction sites of the pSB3K3 vector. Subsequently, the *las* promoter region was replaced by digestion with XbaI and SalI, and the *lux* promoter (iGEM Parts Registry: BBa_R0062) was inserted into the same restriction sites. Randomized promoter libraries were generated analogously using synthesized oligonucleotides in which selected nucleotide positions of the *lux* or *las* promoter were randomized (FASMAC Co., Ltd.), followed by primer extension, digestion with XbaI and SalI, and insertion into the same pSB3K3-P*lux*-GFP or pSB3K3-P*las*-GFP backbone using the corresponding restriction sites. For functional assays, each expression plasmid (or the empty vector pET16b(+)) and a reporter plasmid were co-transformed into *Escherichia coli* DH5α.

### In vivo assays of transcription activation abilities

For individual reporter assays, *E. coli* DH5α cells carrying an expression plasmid and either the reporter plasmid were grown overnight in LB medium supplemented with 50 µg/mL of carbenicillin and 25 µg/mL of kanamycin. Overnight cultures were diluted into fresh LB medium and further grown 1 h at 37°C with shaking. After dilution of fresh culture, protein expression was induced by adding IPTG at a final concentration of 1 mM or 0.33 mM, followed by incubation for 6 h at 37°C. GFP fluorescence and cell density were quantified using either a plate reader (Synergy, Agilent BioTek) or a flow cytometer (CytoFLEX S, Beckman Coulter).

For MPRAs, *E. coli* DH5α cells were similarly co-transformed with a TF expression plasmid and a reporter plasmid. For pooled TF library MPRA, the pooled VAE-designed TF plasmid library was transformed into cells harboring either P*lux* or P*las* reporter plasmid. For the randomized promoter-library MPRA, the promoter-GFP library was transformed into cells harboring an individual TF expression plasmid (LuxR, LasR, 20L, or 22L). In both cases, transformant colonies were collected from agar plates and resuspended into fresh LB medium. In pooled TF library MPRA, resulting suspension was grown 1 h at 37°C with shaking, induced by adding IPTG at a final concentration of 1 mM, followed by incubation for 6 h at 37°C. In the randomized promoter-library MPRA, resulting suspension was induced by adding IPTG at a final concentration of 1 mM, followed by incubation for 10 h at 15°C and further 7.5 h at 37°C. After induction, cells were diluted and subjected to fluorescence-activated cell sorting (FACS). Cells were sorted based on GFP-A fluorescence intensity using a BD FACSAria Fusion Cell Sorter (BD Biosciences, United States) equipped with a 100-µm nozzle. Sorting was performed using the ‘Purity’ precision mode. The GFP-A histogram was partitioned into 8 consecutive gates (bins), and a target of 10,000 cells from each bin were collected into individual wells of a 96-well plate when sufficient events were available, with fewer cells collected for sparsely populated high-fluorescence bins (**Supplementary Fig. S2 and S6**). Each well contained 1/10-strength LB medium supplemented with 50 µg/mL carbenicillin and 25 µg/mL kanamycin. Cells collected from each bin were pelleted, resuspended in 18 µL of Milli-Q water and lysed by heating at 94°C for 5 min. The variable region derived from plasmid DNA (TF coding sequence for the TF library; promoter insert for the promoter library) was amplified by PCR to attach adapter sequences using primers compatible with adapters. To maintain the original population’s profile, the resulting PCR products from each bin were pooled according to the cell distribution ratio observed in the FACS histogram. The pooled products were sequenced using an Illumina NextSeq 1000 Sequencing System, with sequencing services provided by Bioengineering Lab. Co., Ltd. Sequence counts were obtained by exact-match parsing, and enrichment-based activity scores were calculated for each variant by comparing its representation across fluorescence bins.

### Statistical analysis of promoter library activities

To assess whether activity distributions differed significantly among transcription factors, both parametric and nonparametric statistical tests were performed. One-way analysis of variance (ANOVA) was applied to raw activity values, followed by the Kruskal-Wallis test as a distribution-free alternative. Post hoc pairwise comparisons were conducted using Dunn’s test with Bonferroni correction. Homogeneity of variances was assessed using Levene’s test.

For constructing feature matrix *X*, each sequence was encoded using one-hot encoding at the variable positions. For each variable position, four binary features corresponding to nucleotides A, C, G, and T were generated. The response matrix *Y* consisted of activity values for LuxR, LasR, 20L, and 22L. Raw activity values were transformed to reduce skewness. The transformed values were then standardized (zero mean and unit variance) independently for each transcription factor prior to multivariate analysis. Partial least squares (PLS) regression was performed to model the relationship between sequence features and transcriptional activity. PLS was chosen to extract latent components of the sequence feature space that maximize covariance with the activity measurements. The one-hot encoded sequence matrix *X* was used as the predictor matrix, and the standardized activity matrix *Y* was used as the response matrix. Two PLS components were retained for analysis. Model fitting was performed using the PLSRegression implementation from scikit-learn. PLS regression coefficients were extracted to quantify the contribution of each nucleotide at each variable position to transcriptional activity. Coefficients were reshaped into matrices representing nucleotide preferences (A, C, G, T) at each variable position.

## Data availability

All custom code used for VAE model training and latent-space sampling and datasets curated for training the VAE models are available at GitHub (kigalab/VAE-hybridTF). Raw sequencing data from the pooled transcription factor library sort-seq experiments and randomized promoter-library sort-seq experiments have been deposited in Zenodo (10.5281/zenodo.18798256). Additional data supporting the findings of this study are available at Supplementary Information.

## Acknowledgments

We thank R. Yoshida, K. Takahashi, Y. Okada and T. Ueda for critical discussion of the manuscript and valuable suggestions; and the members of the Laboratory of Synthetic Biology (the D. Kiga group) for their support. In particular, we are grateful to H. Shimokawa for assistance with model construction and coding, and to R. Sekine, Y. Takizawa, S. Nishimura and A. Sasaki for providing materials. This work was financed by the JSPS KAKENHI (Grant numbers 19H00985, 21K19831, 21H05228, 22H04892, 23H04431, 24K03036, and 25K22848) to D.K., by JST CREST (Grant number JPMJCR21N4) to D.K. including AIP challenge program to S.L.O. and K.U., and by NINS Astrobiology Center program research (Grant numbers AB041011, AB0602, and AB0712) to D.K.

## CRediT Author contributions

S.L.O.: conceptualization, data curation, investigation, formal analysis, methodology, funding acquisition, software, visualization, writing – original draft, review and editing. A.M.: formal analysis, investigation, methodology, software, visualization, writing – original draft, review and editing. M.A.: formal analysis, investigation, data curation, writing – original draft, review and editing. K.U.: investigation, funding acquisition. K.M.: resources. K.O.: supervision, writing – review and editing. D.K.: conceptualization, formal analysis, methodology, funding acquisition, project administration, resources, supervision, writing - review and editing.

## Competing interests

The authors declare no competing interests.

## Notes

### Competing Interest Statement

The authors have declared no competing interest.

### Summary of Updates

The manuscript has been revised with minor updates to the text, figures, and supplementary information.

