## Supplementary Information for "Generative AI-based design of hybrid transcriptional activator proteins with new DNA-binding specificity"

### Title

* Correspondence to:

### Supplementary Methods

#### *Phylogenetic analysis and reconstruction of ancestral sequences*

Ancestral sequences of the LuxR-family DNA-binding domains were inferred from the same curated MSA used for VAE training. Maximum-likelihood phylogenetic trees were generated using RAxML-NG with the LG+G substitution model[1]. To account for stochastic variation arising from tree-search initialization, three independent trees were constructed by varying the random seed. For each of the three trees, ancestral state reconstruction was performed with --ancestral module of RAxML-NG, producing three independent ancestral sequence predictions.

#### *Detailed Protocol for the Model building and evaluation*

To identify the optimal hyperparameter range where the VAE effectively captures the statistical properties of the LuxR-family DNA-binding domains, we conducted a two-phase grid search. In the first phase, a broad scan was performed across the following parameter space:

(i) the dimension of the latent space: 2, 4, 8, 16, 32, 64, or 128

(ii) the number of units in hidden layers: two layers of 512, 256, or 128

(iii) the β coefficient for the KLD term: 10, 0.1, $10^{-3}$, $10^{-5}$, or $10^{-7}$

(iv) random seeds: 0, 1, or 42

Following the broad scan in the first phase (**Supplementary Fig. S7**), we narrowed the hyperparameter range to focus on the regions that demonstrated high generative fidelity. To ensure the reproducibility of the model performance and to account for the stochastic nature of the training process, the set of random seeds was expanded. Specifically, the second-phase grid search was conducted with the following refined parameters:

(i) the dimension of the latent space: 8, 16, or 32

(ii) the number of units in hidden layers: two layers of 512, 256, or 128

(iii) the β coefficient for the KLD term: $10^{-2}$, $10^{-3}$, or $10^{-4}$

(iv) random seeds: 0, 1, 42, 23461234, or 8747562

#### *Construction of pTrc99A-DsRedExpress2 backbone*

pDsRed-Express2 (Clontech) was digested with NcoI and XbaI, and the DsRedExpress2 coding sequence was inserted into pTrc99A. The resulting plasmid was further digested with XbaI, followed by blunt-end formation using Klenow fragment and self-ligation to eliminate the XbaI site. To introduce additional restriction sites for flexible cloning, PCR was performed to insert SpeI and EcoRI sites upstream of the *trc* promoter, and an NheI site between the *trc* promoter and the coding sequence. The resulting construct was used as the backbone plasmid pTrc99A-DsRedExpress2.

#### *Library construction*

A pool of unique oligonucleotides was synthesized by Integrated DNA Technologies (United States). To generate double-stranded DNA (dsDNA), a primer extension reaction was performed using PrimeSTAR Max DNA Polymerase (Takara Bio, Japan). The reaction mixture contained the oligonucleotide pool and a specific complementary primer at a 1:1 molar ratio. Thermal cycling was carried out with an initial denaturation at 98°C for 1 min, followed by 30 cycles of 98°C for 10 s, 55°C for 15 s, and 72°C for 3 s. The resulting dsDNA was separated on a 12% polyacrylamide gel (PAGE), and the target bands were excised. DNA was eluted from the gel slices by diffusion in buffer at 37°C overnight with rotation, followed by purification by ethanol precipitation. Purified inserts and the pTrc99A-DsRedExpress2 vector were digested with NheI and HindIII and purified by ethanol precipitation. To prevent vector self-ligation, the digested vector was treated with Shrimp Alkaline Phosphatase (Takara Bio). After agarose gel electrophoresis, the desired vector band was excised and purified using the NucleoSpin Gel and PCR Clean-up kit (MACHEREY-NAGEL, Germany) according to the manufacturer's instructions. Ligation was performed using the DNA Ligation Kit <Mighty Mix> (Takara Bio). The vector (50 ng) and insert were mixed at a 1:10 molar ratio, heated at 65°C for 10 min, and rapidly chilled on ice prior to addition of the ligation mixture. The ligation products were purified and concentrated by ethanol precipitation for desalting. The purified ligation products were electroporated into *E. coli* DH5α electrocompetent cells (Takara Bio) using a MicroPulser Electroporator (Bio-Rad, United States). Immediately after pulsing, 1 mL of SOC medium was added, and cells were incubated at 37°C for 30 min for recovery before plating on LB agar containing ampicillin. Plates were incubated overnight at 37°C. Colonies were collected from the plates and resuspended in LB medium, and plasmid DNA was extracted from the resulting bacterial suspension. Approximately 1,500 (P*las*N5), 6,000 (P*lux*N6), and 3,000 (120-pool library) transformant colonies were obtained.

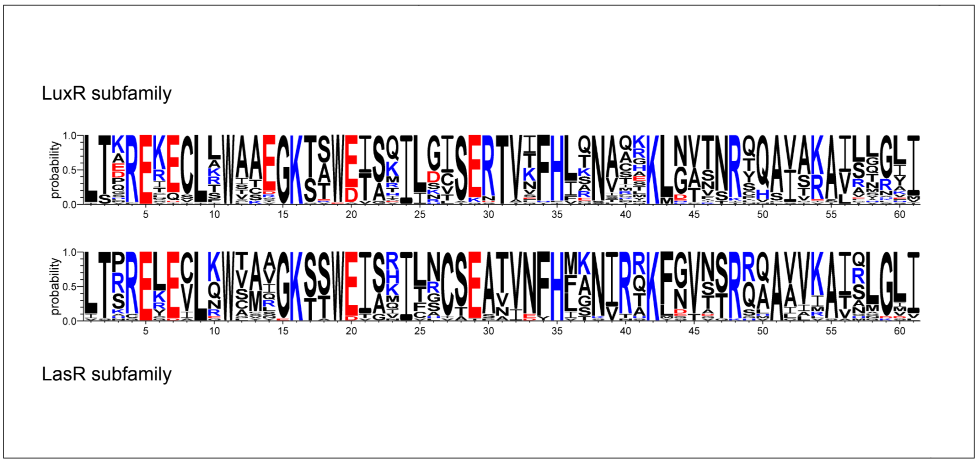


**Supplementary Fig. S1 | Sequence logos of LuxR and LasR subfamily DNA-binding domains**

For each subfamily, sequences were extracted from the curated LuxR-family dataset by using wild-type LuxR or LasR as a BLAST query and selecting hits with ≥60% pairwise identity. Redundant sequences were removed by CD-HIT clustering at 90% identity, and the remaining representatives were aligned using MAFFT. Sequence logos were generated from the resulting multiple sequence alignments to visualize subfamily-specific residue conservation patterns.

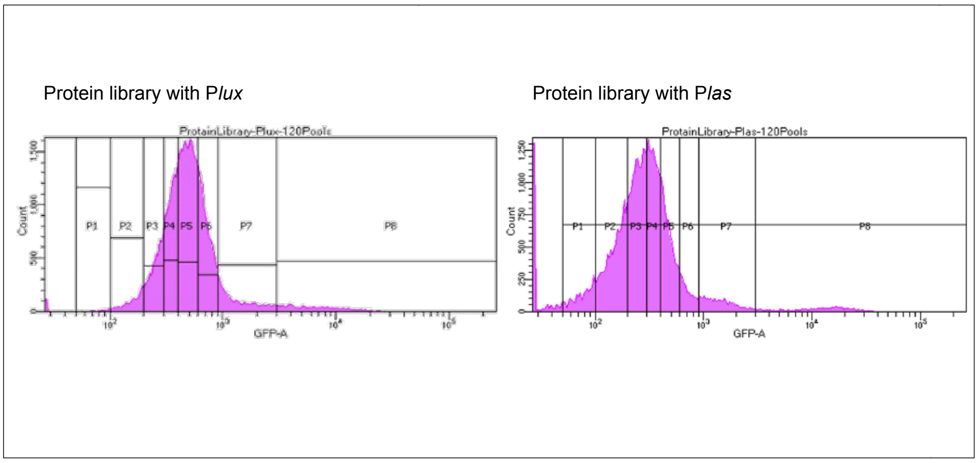


**Supplementary Fig. S2 | FACS fluorescence distributions for sort-seq of the protein library**

Histograms show single-cell GFP fluorescence measured by FACS prior to sorting for *E. coli* cells expressing the pooled library of 120 VAE-designed transcription factor variants together with the native *lux* or *las* promoter reporter. Vertical lines indicate the fluorescence bin boundaries used to define sorting bins. These bins were used for subsequent recovery of plasmid DNA and deep sequencing to calculate enrichment-based activity scores for each variant.

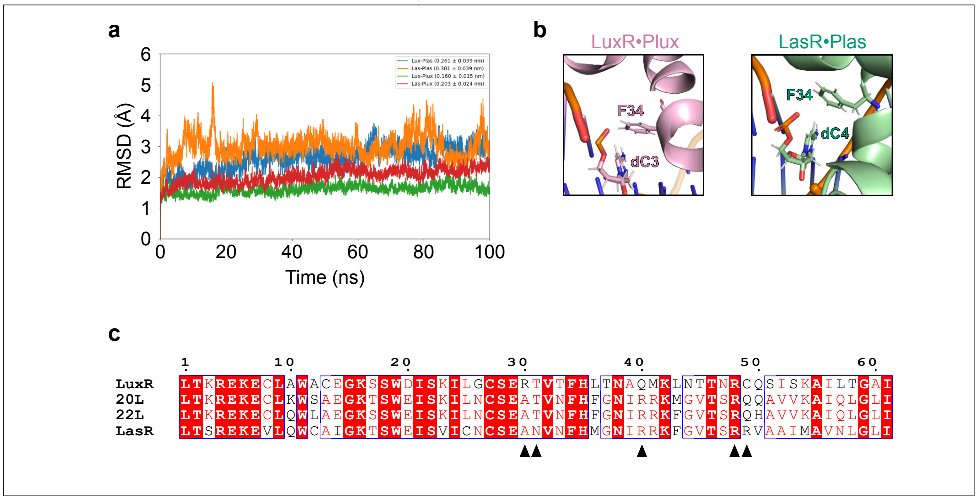


**Supplementary Fig. S3 | Molecular dynamics simulations and sequence features underlying promoter recognition (related to Fig. 3d)**

**a,** Root-mean-square deviation (RMSD) trajectories from 100 ns molecular dynamics (MD) simulations of four TF-DNA complexes: LuxR•Plux, LuxR•Plas, LasR•Plux, and LasR•Plas. RMSD values were calculated for each complex relative to its respective starting structure. **b,** Representative MD-refined structures highlighting interactions between Phe34 and the cognate DNA bases. Enlarged views of the LuxR•Plux (left) and LasR•Plas (right) complexes after 100 ns of MD simulation are shown, focusing on the local environment around Phe34. The aromatic side chain of Phe34 and the corresponding DNA bases are displayed to illustrate a T-shaped π-π stacking interaction. **c,** Multiple sequence alignment of LuxR, LasR, and VAE-designed variants (20L and 22L). Arrowheads indicate amino acid positions explicitly discussed in the main text with respect to promoter recognition and binding energetics.

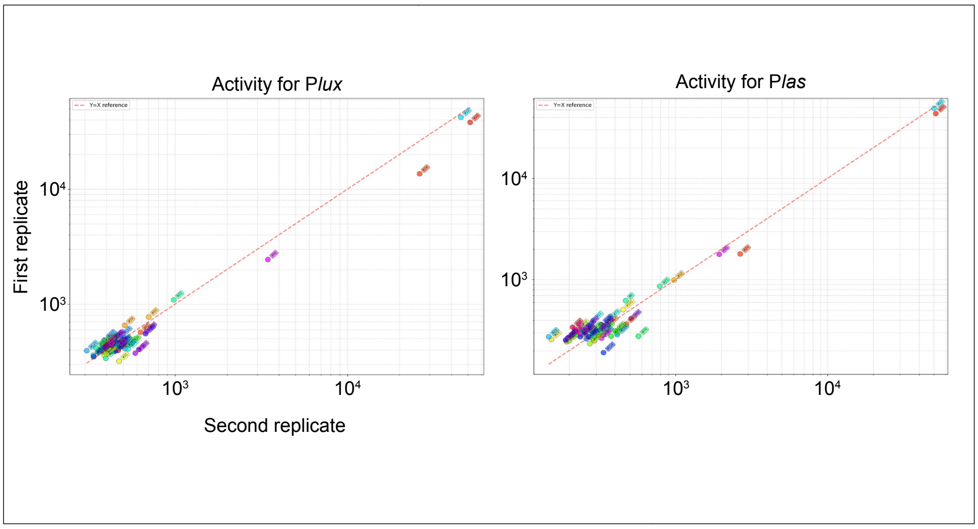


**Supplementary Fig. S4 | Reproducibility of protein-library sort-seq measurements across biological replicates (related to Fig. 3b)**

Scatter plots comparing promoter-activation scores obtained from two independent biological replicate sort-seq experiments for the protein library. For each variant detected in both experiments (n = 57), the geometric mean activity score from the first replicate (shown in Fig. 3b) is plotted on the y-axis and that from the second replicate on the x-axis. The second replicate yielded fewer sequencing reads than the first. Therefore, only variants with at least 20 total sequence counts in both experiments were included in this analysis, resulting in a subset of the variants shown in Fig. 3b. Panels show results for the *lux* promoter (left) and *las* promoter (right), respectively. Pearson correlation coefficients (r) are 0.9847 (Plux) and 0.9971 (Plas), respectively. Activity scores were calculated from exact sequence counts across fluorescence bins, as described for Fig. 3b.

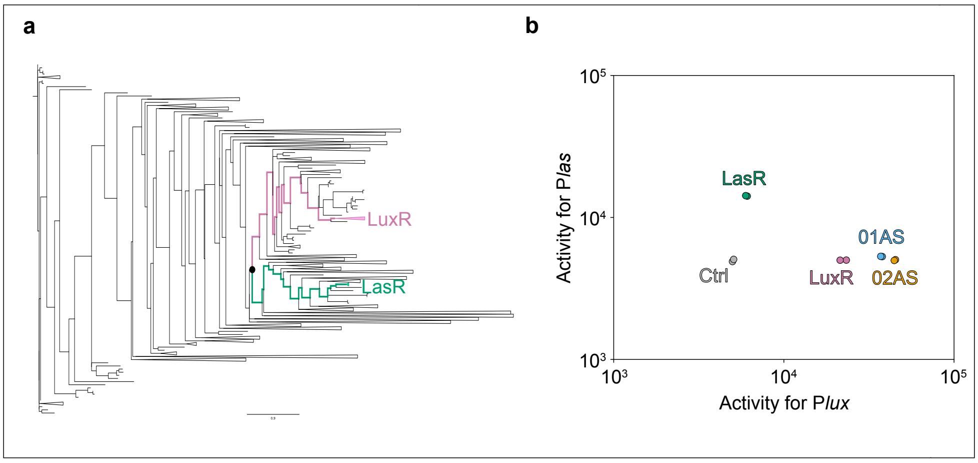


**Supplementary Fig. S5 | Ancestral sequence reconstruction and functional evaluation**

**a,** Maximum-likelihood phylogenetic tree used for ancestral sequence reconstruction (one of three independent reconstructions). Ancestral nodes inferred from each reconstruction are indicated. **b,** Transcriptional activation abilities of the three reconstructed ancestral DBDs toward the *lux* and *las* promoters. Each point represents an individual biological replicate (n = 2). One ancestral variant (03AS) exhibited poor growth after induction and could not be reliably assayed.

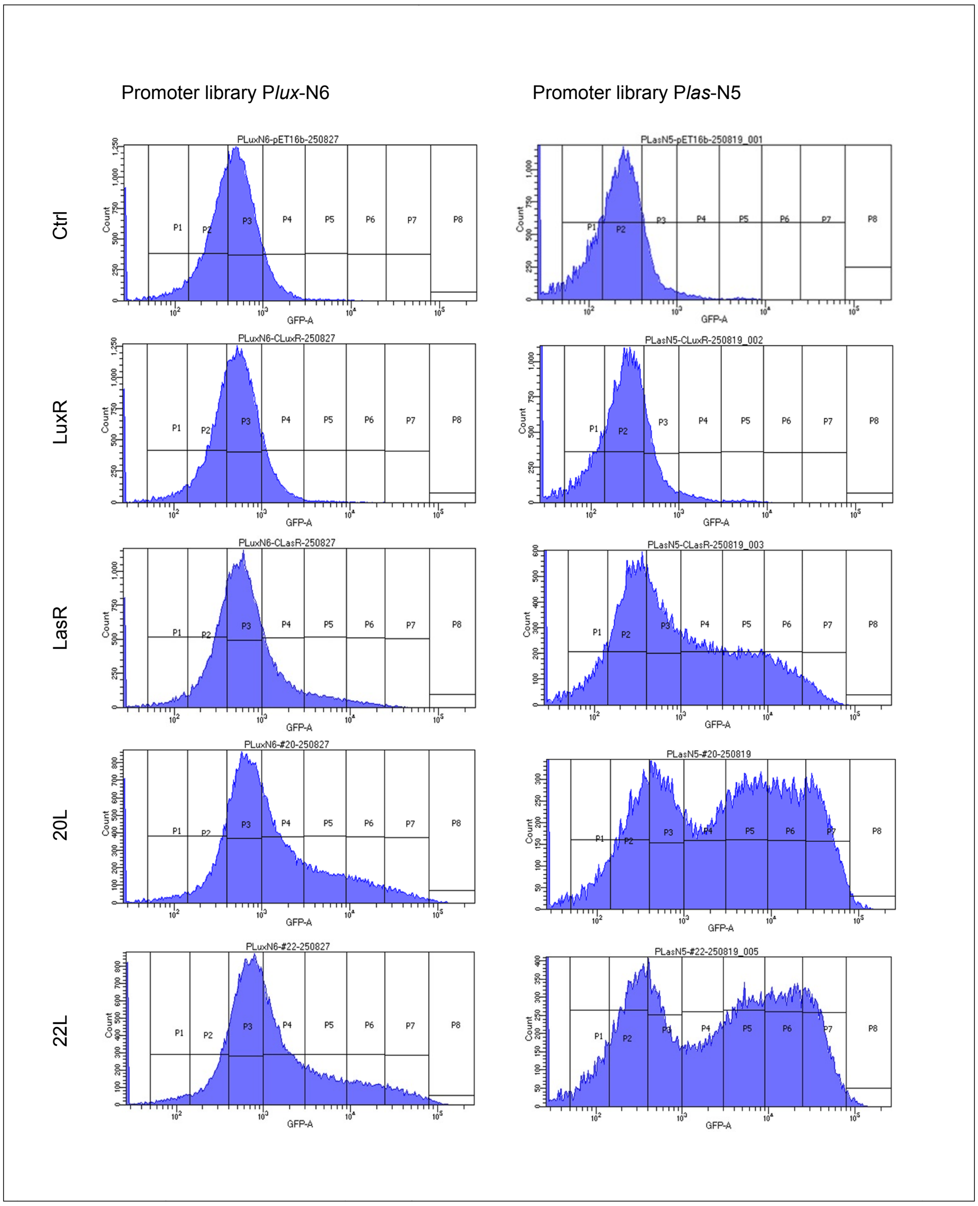


**Supplementary Fig. S6 | FACS fluorescence distributions for sort-seq of the promoter library**

Histograms show single-cell GFP fluorescence measured by FACS prior to sorting for cells expressing wild-type LuxR, wild-type LasR, or VAE-designed variants (20L or 22L) together with the randomized promoter libraries. Vertical lines indicate the fluorescence thresholds used to define sorting bins. These bins were used for subsequent recovery of plasmid DNA and deep sequencing to calculate enrichment-based activity scores.

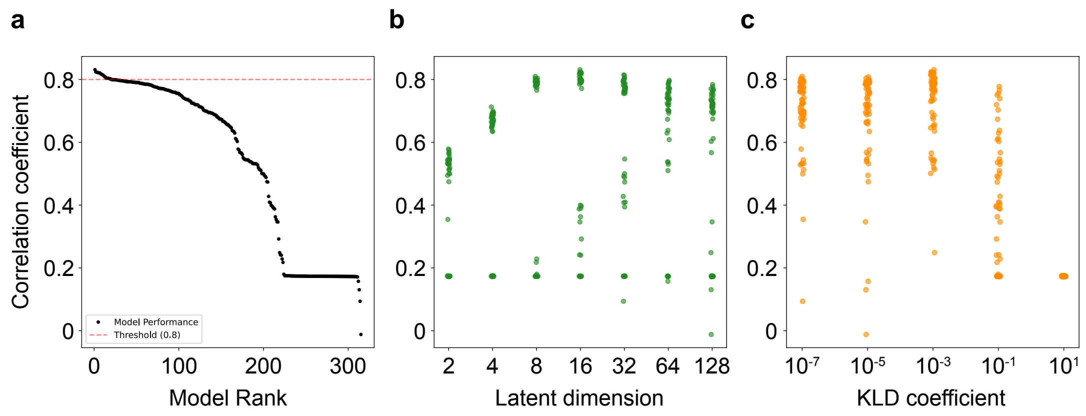


**Supplementary Fig. S7 | Evaluation of VAE models through grid search**

**a,** Distribution of co-mutation correlation coefficients across all models. Models are sorted in descending order based on the correlation coefficient of amino acid pair frequencies calculated from sequences generated via spherical interpolation in the latent space. The horizontal dashed line indicates the threshold of 0.8. **b,** Impact of latent space dimensionality on model performance. Co-mutation correlation coefficients are plotted against the dimension of the latent space. Points are slightly jittered for better visualization. High-performance models were predominantly observed in dimensions of 8, 16, or 32. c, Influence of the KL divergence coefficient on interpolation capability. The relationship between the β coefficient (weight of the KLD term) and the co-mutation correlation coefficient is shown. Optimal performance was achieved at $10^{-3}$.

**Supplementary Table S1 | Performance and hyperparameters of the top 10 VAE models**

| Model ID | Latent Dim. | Hidden layer architecture | KLD coef. | Random seed | Correlation Value |
|---|---|---|---|---|---|
| 94 | 32 | 512-512 | 0.01 | 8747562 | 0.8330 |
| 107 | 32 | 256-256 | 0.01 | 42 | 0.8325 |
| 109 | 32 | 256-256 | 0.01 | 8747562 | 0.8319 |
| 52 | 16 | 512-512 | 0.001 | 42 | 0.8308 |
| 53 | 16 | 512-512 | 0.001 | 23461234 | 0.8297 |
| 69 | 16 | 256-256 | 0.001 | 8747562 | 0.8270 |
| 92 | 32 | 512-512 | 0.01 | 42 | 0.8267 |
| 90 | 32 | 512-512 | 0.01 | 0 | 0.8261 |
| 51 | 16 | 512-512 | 0.001 | 1 | 0.8253 |
| 84 | 16 | 128-128 | 0.001 | 8747562 | 0.8243 |

**Supplementary Table S2 | Plasmids used in this study**

| Name | Description | Source |
|---|---|---|
| pPtet LuxR | Expressing LuxR under *tet* promoter | Sekine *et al*. (2011)[2] |
| pSB6A1 | A plasmid with an ampicillin resistance marker | iGEM Parts |
| pSB6A1-P*tet*-C*luxR* | Expressing the C-terminal region of LuxR under *tet* promoter | This study |
| pDsRed-Express2 | A plasmid coding DsRedExpress2 | Clontech |
| pTrc99A | A plasmid with an ampicillin resistance marker | Amann *et al*. (1988)[3] |
| pTrc99A-DsRedExpress2 | Expressing DsRedExpress2 under *trc* promoter | This study |
| pTrc99A-C*lasR* | Expressing the C-terminal region of LasR under *trc* promoter | This study |
| pSB3K3 | A plasmid with a kanamycin resistance marker | iGEM Parts |
| pSB3K3-P*lux*-GFP | Expressing GFP under *lux* promoter | This study |
| pSB3K3-P*las*-GFP | Expressing GFP under *las* promoter | This study |

184 **Supplementary Table S3 | Primer sequences used in this study**

| Name | Sequence (5’-3’) N: randomized base |
|---|---|
| Ptet-luxR_del2-182_FW | AGGATTTCTACAATGTTAACCAAAAGAGAAAAAGAATGTTTAGCG |
| Ptet-luxR_del2-182_RV | TATTGACTTATTAAACTAGTAGTGCTCAGTATCTCTATCACTG |
| pTrc99A_EcoRISpeI_57_short_FW | ATAACGGAATTCGTTCTGGCAAATATTCTGAAATGAGC |
| pTrc99A_EcoRISpeI_57_short_RV | GATGACTAGTTCGGCGCAAAAAACATTATCC |
| pTrc99A_NheI_short_FW | AGCTTAAAGAAGGAATCGGAGGAGAATTAATGGATAGCACTGAGAACGTCA<br>TCAAG |
| pTrc99A_NheI_short_RV | GCTAGCTTGTTATCCGCTCACAATTCCACACA |
| ClasR_oligo | TAACAAGCTAGCCGCTTGAAATAATTGATAACAACCCCCTAGGCAGCAATTT<br>AATGTTAACGAGTCGTGAAAAAGAAGTCTTGCAGTGGTGTGCGATTGGGAAA<br>ACGTCATGGGAAATCTCGGTTATCTGCAACTGCTCTGAAGCTAATGTAAACTT<br>TCACATGGGAAACATTCGCCGTAAATTCGGCGTGACTTCTCGCCGCGTTGCG<br>GCAATCATGGCTGTGAATCTTGGGCTTATTTAATAAGCTTGGCTGTTTTGGCG<br>GA |
| 20L_oligo | TAACAAGCTAGCAACGAACCAGCAATTGGGGAGCAAAAATGGCGCTTACAA<br>AACGTGAAAAAGAATGTCTGAAGTGGTCGGCGGAGGGCAAAACCTCTTGGG<br>AAATTAGCAAAATTTTGAATTGCAGCGAAGCAACGGTTAATTTTCATTTTGGT<br>AATATCCGTCGCAAGATGGGGGTCACAAGTCGCCAGCAGGCAGTAGTAAAG<br>GCTATCCAATTAGGCTTAATCTAATAAAAGCTTGGCTGTTTTGGCGGA |
| 22L_oligo | TAACAAGCTAGCGACCCGGGTATACGGAGGTCATTGATGGCGTTAACCAAAC<br>GTGAAAAAGAATGCTTGCAGTGGCTTGCGGAAGGCAAAAGCTCGTGGGAAA<br>TTTCAAAGATTCTTAATTGTTCCGAGGCAACAGTCAACTTCCACTTCGGAAAC<br>ATCCGTCGTAAATTCGGTGTAACTTCGCGCCAACATGCCGTTGTGAAAGCAA<br>TCCAGCTTGGCCTGATTTAATAAAAGCTTGGCTGTTTTGGCGGA |
| PluxN6_oligo | GCTTCTAGAGTTCGAGCCTAGCAAGGGTCCGGGTTCACACCTNNNGGATCGN<br>NNAGGTTTACGCAAGAAAATGGTTTGTTATAGTCGAATAAATACTAGAGGTC<br>GACTGA |
| PlasN5_oligo | GCTTCTAGAGTTCGAGCCTAGCAAGGGTCCGGGTTCACCGAAACCTNNNNNA<br>TTTGCTAGTTATAAAATTATGAAATTTGCGTAAATTCTTCATACTAGAGGTCG<br>ACTGA |
| 120Lib_extension | TCCGCCAAAACAGCCAAGCTT |
| PluxN6_extension | TCAGTCGACCTCTAGTATTTATTCGACTATAACAAACC |
| PlasN5_extension | TCAGTCGACCTCTAGTATGAAGAATTTACGCAAATTTC |
| 120Lib_NGS_FW | ACACTCTTTCCCTACACGACGCTCTTCCGATCT[barcode]ttgacaattaatcatccggctc<br>barcode:<br>Plas_P1: CGTCGCAC<br>Plas_P2: CGTCGTCA<br>Plas_P3: CGTACGCA<br>Plas_P4: CGTACGCT<br>Plas_P5: CGAGTGCC<br>Plas_P6: CGATCGCA<br>Plas_P7: ACGCGTCA<br>Plas_P8: CGCTGCGC<br>Plux_P2: CGCGATCA<br>Plux_P3: CGCTGGCA<br>Plux_P4: CGCTGACA<br>Plux_P5: CGCTAGCA<br>Plux_P6: CGCAGTCA<br>Plux_P7: CGCATGCA<br>Plux_P8: CGTGAGCC |
| 120Lib_NGS_RV | GTGACTGGAGTTCAGACGTGTGCTCTTCCGATCT[barcode]atccgccaaaacagccaag<br>barcode:<br>Plas_P1: GTCGTACG<br>Plas_P2: GTACGCGA<br>Plas_P3: GTACGTCG<br>Plas_P4: CGCGTGAG<br>Plas_P5: GTGTGAGA<br>Plas_P6: GTGCGACG<br>Plas_P7: GTCGTACA |

| | |
|---|---|
| | Plas_P8: GTCGATCG<br>Plux_P2: GCATGCAG<br>Plux_P3: GTGCGATG<br>Plux_P4: GTGCGAGA<br>Plux_P5: GTGAGCGA<br>Plux_P6: GTCGCGTG<br>Plux_P7: GTCGCAGG<br>Plux_P8: GTCGTCGA |
| PromLib_NGS_FW | ACACTCTTTCCCTACACGACGCTCTTCCGATCT[barcode]tttctggaattcgcggccgctt<br>barcode:<br>P1: CGCGATCG<br>P2: ACGCGTCG<br>P3: AGCGCATG<br>P4: CGACGCTG<br>P5: CGCTAGCG<br>P6: CGCATGCG<br>P7: CGATCGCG<br>P8: CGTCGCGA |
| PromLib_NGS_RV | GTGACTGGAGTTCAGACGTGTGCTCTTCCGATCT[barcode]tccgacaggatccagtcg<br>barcode:<br>P1: GCATGCTG<br>P2: CGCGAGTG<br>P3: GATCGCGA<br>P4: GAGCGTGA<br>P5: GTGAGCGC<br>P6: GTCGCAGC<br>P7: GAGCGTCG<br>P8: GTCGTACG |
